# Neural entrainment to speech in theta range is affected by language properties but not by the native language of the listeners

**DOI:** 10.1101/2023.07.11.548540

**Authors:** Ege Ekin Özer, Silvana Silva Pereira, Nuria Sebastian-Galles

## Abstract

A universal speech rhythm around 5 Hz, corresponding to syllable beats, is captured by neural oscillations in the human brain. However, there is significant variability in syllabic complexity across languages: some languages allow only simple syllables, while some allow more variation (a variation related to linguistic rhythm). Behavioral evidence suggests that humans show different patterns of speech segmentation depending on the linguistic rhythm of their native language. Here, we tested if the entrainment of neural oscillations in the theta range (3–8 Hz) to sentences of languages representative of different linguistic rhythms depends on participants’ native language rhythm or rather reflects language-specific rhythmic properties. We recorded EEG in two groups of participants: native speakers of English (stress-timed, Experiment 1) and native speakers of Spanish (syllable-timed, Experiment 2). Both groups listened to *saltanaj* resynthesized sentences from English (stress-timed), Spanish (syllable-timed) and Japanese (mora-timed), a procedure that removes comprehension but keeps language-specific phonological properties. Phase locking value between sentence envelopes and EEG showed the same pattern regardless of participants’ native language: lowest for English, intermediate for Spanish, and highest for Japanese. Our results suggest that entrainment to speech in the theta frequency range is sensitive to differences in variation in syllabic complexity across languages.

## Introduction

Neural models of speech perception propose that the brain segments speech into syllable-like chunks through entrainment in the theta frequency range (3–8 Hz) and assume that entrainment in this range is signal-driven (Giraud and Poeppel, 2012; Poeppel and Assaneo, 2020; Rimmele et al., 2018). Moreover, entrainment is associated with increased intelligibility and comprehension (Peelle et al., 2013; Bosker and Ghitza, 2018). Phonological knowledge is considered responsible for bottom-up entrainment (Bourguignon et al., 2013). However, phonological information is not isomorphic to acoustic information. There is abundant evidence that the perception of acoustic information depends on the listeners’ native language (Sebastian-Gallés et al., 2006; Luo et al., 2007; Li et al., 2021) and that language-specific phonological properties, in particular linguistic rhythm, influence the way listeners parse the speech signal.

World languages have been classified into three rhythmic classes: stress-timed (English or Arabic), syllable-timed (Spanish or Yoruba) and mora-timed (Japanese or Tamil), assuming isochrony at the level of feet, syllables, and morae, respectively (Pike, 1945; Abercrombie, 1967). Recently, linguistic rhythm has been associated with syllabic complexity (Ramus et al., 1999; Langus et al., 2017): complex syllabic structures, composed of much higher ratio of consonants (C) than vowels (V), are only found in stress-timed languages (such as the English word “splint” CCCVCC), intermediate structures in syllable-timed languages, while mora-timed languages only allow simple structures (mostly CV and V). Consequently, syllable rates are less regular in stress-timed languages, intermediate in syllable-timed ones, and more regular in mora-timed languages (Pellegrino et al., 2011).

There is abundant evidence that linguistic rhythm affects speech segmentation. In a seminal study by Mehler et al. (1981), native speakers of French listened to words and were instructed to respond as soon as they heard predetermined target sequences. Response times were faster when the target coincided with the first syllable of the word. However, Cutler et al. (1986) found no trace of such syllabic processing in English speakers, concluding that they do not segment speech based on syllables. A second line of research investigated the effects of cross-language adaptation to highly compressed speech (Pallier et al., 1998), showing better performance when adaptation sentences belonged to the same linguistic rhythm as participants’ native language. These studies have been taken to indicate that native linguistic rhythm affects speech segmentation.

Experiments on neural entrainment to speech using M/EEG (magneto/electroencephalography) provide mixed results about native phonological specialization. On the one hand, Peña and Melloni (2012) found higher spectral power in the gamma band to sentences in participants’ native language, and Pérez et al. (2015) found higher spectral power in theta to native sentences. On the other hand, Etard and Reichenbach (2019) reported that native speakers of English showed similar cross-correlation between Dutch and English in the theta band. Ding et al. (2016) reported comparable spectral peaks in the theta range when native speakers of English and those of Mandarin listened to Mandarin sentences (Mandarin has been described as a syllable-timed language).

The present research investigates the impact of cross-language rhythmic information on neural entrainment. With this aim, we recorded participants’ EEG while listening to sentences from stress-, syllable-, and mora-timed languages (English, Spanish, and Japanese, respectively). To remove the confounding effects of familiarity and comprehension, we used *saltanaj* resynthesized sentences (Ramus and Mehler, 1999). We conducted two experiments (Experiment 1: native speakers of English, Experiment 2: native speakers of Spanish) with the same set of sentences from each language. To capture entrainment at the syllable level, we computed phase locking value (PLV) between EEG signals and speech envelopes in the theta range. If speech segmentation is tuned to the listeners’ language rhythm, neural entrainment should be higher for the sentences belonging to the listeners’ native language. However, if neural entrainment is driven by the stimulus rhythm, all participants should show the same pattern of entrainment, regardless of their native language.

## Methods

### Participants

Twenty-four native speakers of English (aged 23.71 ± 4.07) participated in Experiment 1 (M = 8, F = 16). Twenty-four native speakers of Spanish (aged 23.17 ± 4.74) participated in Experiment 2 (M = 9, F = 15). Participants’ age range was from 18 to 35 years. The participants were assigned to the groups according to which language they grew up speaking to their parents or guardians. Participants that took part in our experiments were contacted mainly through the participant database of the Center for Brain and Cognition. Some participants were contacted through flyers distributed on campus and through the advertising of our experiment on social media. Participants were eligible if they were not proficient in the non-native languages presented, depending on their native language: In Experiment 1, native English participants did not have a broad knowledge of Spanish and Japanese, and in Experiment 2, native Spanish participants did not have a broad knowledge of English and Japanese. Secondly, as the experiment was run in Barcelona, Spain, English-speaking participants were selected only if they had spent less than six months in a Spanish-speaking country. Sample sizes were decided in accordance with other studies studying neural entrainment to speech (e.g., Peña and Melloni 2012; Pérez et al. 2015).

All participants reported right-handedness and no hearing problems. They signed an informed consent before starting the experiment and were paid 20 euros for their participation. The experimental procedures were approved by the Drug Research Ethical Committee (CEIm) of the IMIM Parc de Salut Mar, reference 2020/9080/I.

### Stimuli

We used 100 *saltanaj* resynthesized sentences in Dutch and English (stress-timed languages), Spanish and Catalan (syllable-timed languages), and Japanese (mora-timed language) with 20 sentences in each language (Table 1). The sentences had already been used in previous research (Ramus and Mehler 1999; Johnson et al. 2003; F. Ramus, personal communication). The resynthesis method was applied to 20 different declarative sentences, recorded by four female native speakers of each language (each speaker contributed with 5 sentences). The *saltanaj* method is conducted on Bliss (Mertus, 1989) and MBROLA softwares (Dutoit et al., 1996), as well as custom-made code: first, the pitch and duration of the utterances are measured, then the phonemes are identified. The phoneme durations and pitch curves from the original sentences are preserved (Ramus, 2002). Then, the phonemes are replaced by their linguistic equivalents (i.e., phonemes that are common to most languages): all fricatives are replaced with /s/, vowels with /a/, liquids with /l/, stop consonants with /t/, nasals with /n/, and glides with /j/ (Ramus and Mehler, 1999). For example, the English sentence “The next local elections will take place during the winter” would be transformed into “sa natst latl alatsans jal taat tlaas tjalan sa janta.” More examples of *saltanaj* resynthesized sentences can be found online (Speech resynthesis).

**Table 1:**
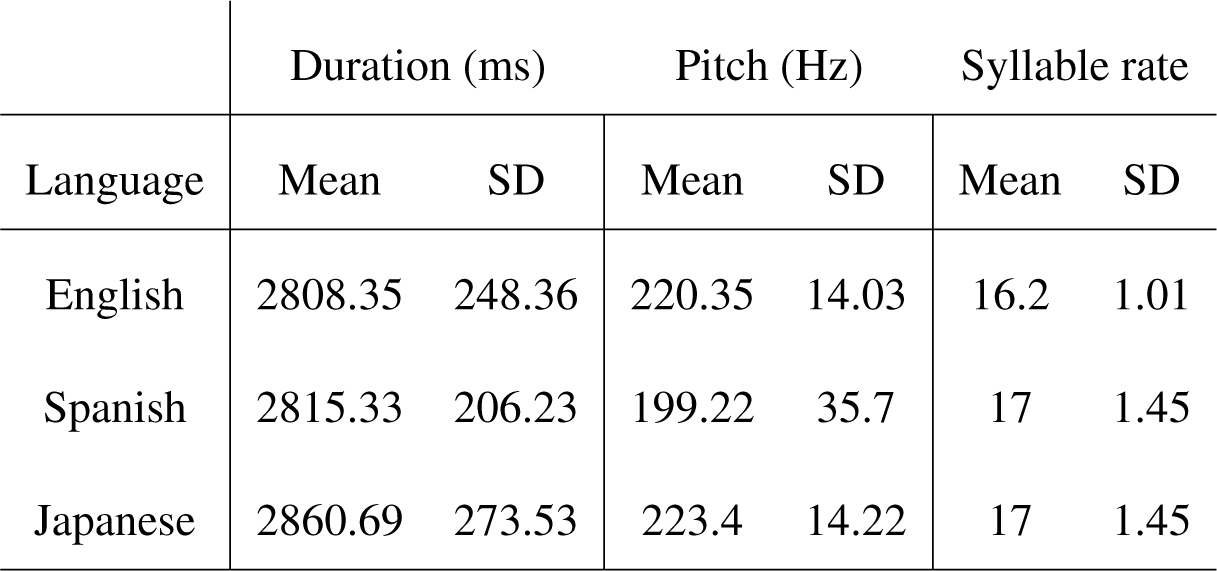
Summary of acoustic and phonological properties of the stimuli used in the experimental blocks.

| Language | Duration (ms) |  | Pitch (Hz) |  | Syllable rate |  |
| --- | --- | --- | --- | --- | --- | --- |
|  | Mean | SD | Mean | SD | Mean | SD |
| English | 2808.35 | 248.36 | 220.35 | 14.03 | 16.2 | 1.01 |
| Spanish | 2815.33 | 206.23 | 199.22 | 35.7 | 17 | 1.45 |
| Japanese | 2860.69 | 273.53 | 223.4 | 14.22 | 17 | 1.45 |

### Procedure

The experiment took place at the Center for Brain and Cognition, Pompeu Fabra University in Barcelona, Spain. Participants sat comfortably on an armchair in an electrically shielded and sound-attenuated room. EEG caps were placed on their heads (see EEG Data Acquisition and Preprocessing for details). The stimuli were presented via Psychtoolbox (Brainard and Vision, 1997) with two Creative T-20 loudspeakers on both sides of the computer screen at a comfortable distance in front of the participants (approximately 120 centimeters).

At the beginning of the experiment, participants read the instructions in their native language presented on the computer screen. Then, participants were informed that they would listen to a series of sentences from an unknown language, and that they would be given a behavioral task to perform for certain sentences. The experiment consisted of four blocks, starting with one training block followed by three experimental blocks: in each block, resynthesized sentences from a single language were presented. After every block, a piece of instrumental music was played for 20 seconds (*Mariage d’amour* performed by Paul de Senneville, *Ayrılık* and *Fikrimin ince gülü* performed by Farid Farjad), to disrupt the sustained neural oscillations previously entrained to speech (van Bree et al., 2021). When the music stopped, 10 seconds of silence ensued and participants were allowed to move on to the next block at their own pace.

Each block consisted of 40 trials: we achieved this by presenting 20 resynthesized sentences twice in each block while ensuring that the same sentence was never presented consecutively. To direct participants’ attention to the sentence onset and indicate that they should be still, every trial started with the appearance of an asterisk in the center of the screen (Figure 1A). Three hundred milliseconds after the appearance of the asterisk, a resynthesized sentence was played. Three hundred milliseconds after the sentence onset, the asterisk disappeared but participants were instructed to keep fixating the asterisk’s location without moving or blinking. Approximately two seconds after each sentence, participants were shown a blue cross for two seconds, indicating that they could blink during that period, as they had previously been instructed. Out of 40 trials, 10 were randomly selected to include a behavioral task aimed to keep participants engaged and attentive to the experiment, particularly on the syllables in the sentences (Figure 1B). The behavioral task appeared approximately two seconds after the presentation of the resynthesized sentences, where the interval was calculated by adding 1 to a number uniformly distributed between 0 and 1 and generated with MatLab’s function *rand*. Participants were presented with two syllables on the screen (out of the 4 options “sa”, “la”, “na”, “ta”), and were asked to estimate which one had occurred more frequently in the last sentence they had heard. We did not include “ja” as a candidate syllable because we did not want the pronunciation of the letter “j” to interfere with the phoneme /j/ heard during the experiment. Using the index and middle fingers of their right hand, participants were instructed to press “1” on the number pad of the keyboard to select the syllable on the left-hand side of the screen and “2” to select the syllable on the right-hand side. Participants were further instructed to respond as quickly as possible with a maximum response time of 3 seconds. Response options remained on the screen until the participant responded or the 3-second period was reached. To ensure speed in response, and to avoid noise in the EEG, we asked participants to always keep their index and middle fingers on the “1” and “2” keys on the number pad throughout the experiment. The first five trials of the training block did not include this behavioral task to allow participants to familiarize themselves with the stimuli.

**Figure 1:**
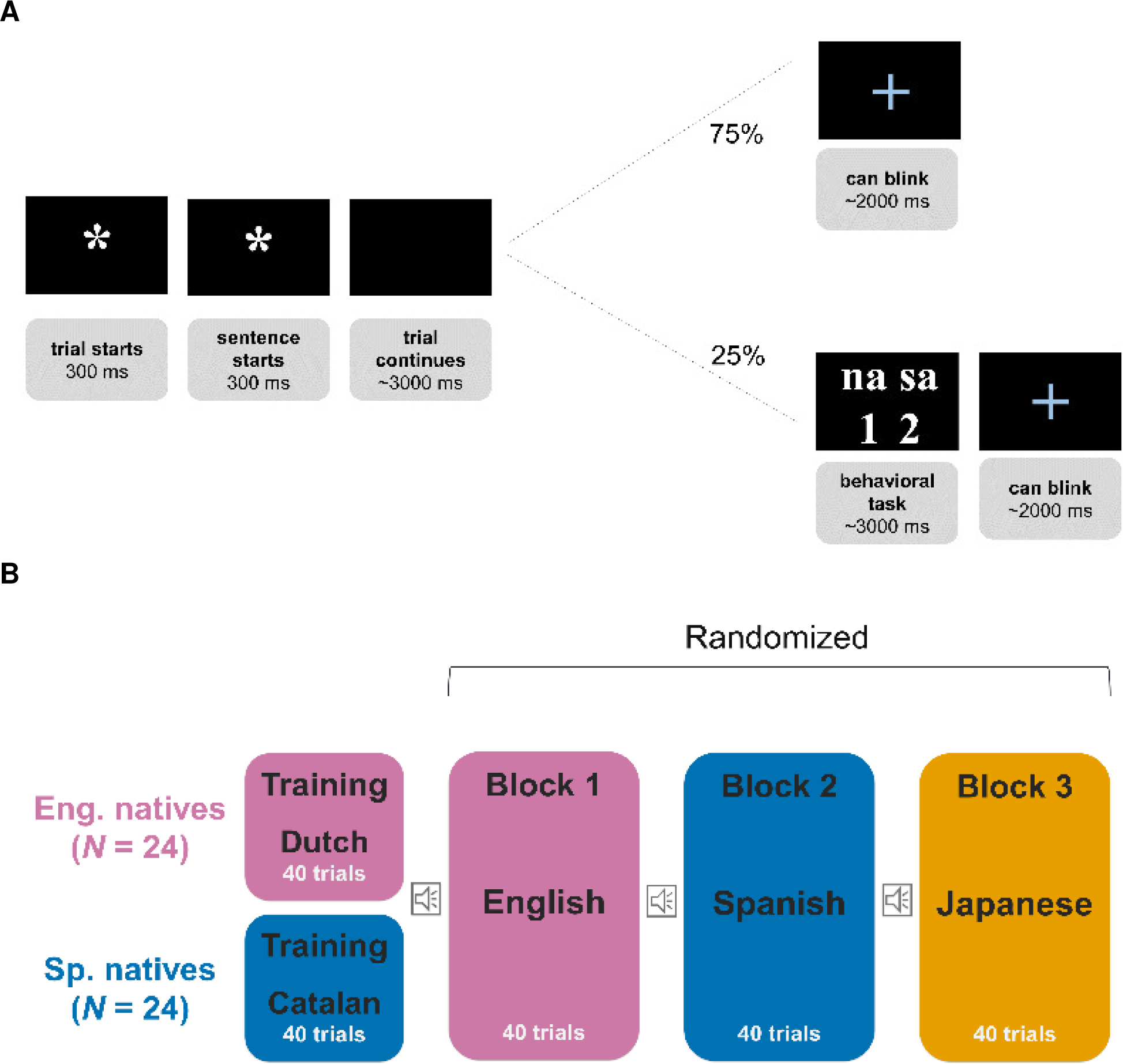
Experimental procedure. A) Trial sequence with and without the behavioral response. Appearance of the asterisk on the screen indicates the start of the trial, participants are told to fixate on the asterisk for the duration of the trial. Three hundred milliseconds after the appearance of the asterisk, the sentence starts playing. Three hundred milliseconds after sentence onset, the asterisk disappears, but the participants are instructed to keep fixating the asterisk’s location. Seventy-five percent of the trials in each block do not require a behavioral response. In such trials, participants continue to listen to the sentence until, once finished, a blue cross appears on the screen, indicating that participants can blink. Twenty-five percent of the trials in each block require a behavioral response. These trials included a behavioral task following the end of the sentence and otherwise followed a sequence identical to those without. Two syllables appeared on the screen with the corresponding keys participants had to press to respond. The size of the elements in this figure (asterisk, text, and the blue cross) is adjusted for visualization purposes. B) Illustration of the experimental procedure showing the order of conditions from different languages. Participants were first presented with the training block, in which they listened to resynthesized sentences from a language rhythmically similar to their native one. They were then presented with the experimental blocks, with resynthesized sentences from stress-, syllable-, and mora-timed languages, in randomized order. In each block, participants were presented with sentences from only one language.

In the training block, participants listened to resynthesized sentences from a non-native language with the same linguistic rhythm as their native language. In the following three blocks, participants listened to resynthesized sentences from a stress-timed language, a syllable-timed language, and a mora-timed language, in counterbalanced order. In Experiment 1, native English participants listened to Dutch in the training block, English in the stress-timed language block, Spanish in the syllable-timed language block, and Japanese in the mora-timed language block. In Experiment 2, native Spanish participants listened to Catalan in the training block, English in the stress-timed language block, Spanish in the syllable-timed language block, and Japanese in the mora-timed language block (Figure 1B).

### EEG Data Acquisition and Preprocessing

EEG was recorded at 1000 Hz with online high-pass filtering over 0.1 Hz via a 64-channel Acticap on BrainVision Recorder and BrainAmp amplifier (Recorder, Vers. 1.22.0001). Channel distribution was in accordance with the International 10/20 system. Event marker triggers were sent via MatLab during the recording. One channel was placed on the nose as the online reference, as well as two outer channels on the left and right mastoid bones to be averaged offline. Finally, two channels were placed horizontally and vertically next to and under the right eye respectively, in order to record eye movements and blinks. Impedances were kept below 10 kΩ.

Data preprocessing was conducted on MatLab using the Fieldtrip toolbox (Oostenveld et al., 2011). Independent component analysis was conducted on the continuous EEG signals to identify and eliminate heart and eye movement artifacts. Then, continuous data were segmented into trials by delimiting one second before and 3.5 seconds after the sentence onset. Power line frequency was removed from each trial and the signals were re-referenced to the mastoidal channels. Noisy channels and trials were visually inspected and eliminated. Channels that were eliminated or deemed noisy were interpolated using spherical spline algorithms to the neighboring channels employed by triangulation. The signals were filtered using a minimum order bandpass FIR filter with Kaiser window between 3 and 8 Hz, with 60 dB attenuation at 2 Hz and 9 Hz, respectively. To minimize the effects of volume conduction with EEG data, we applied a surface Laplacian filter on the signals. Electrode distances were computed using the 3D positions using spherical spline method, and the EEG potential distribution of each channel was recalculated.

Due to an experimenter error, one of the 20 English sentences was saved with two different names. This resulted in all participants in Experiment 1 and four participants in Experiment 2 listening to this sentence four times, instead of only twice. To avoid any potential effects of repetition in our results, we eliminated the trials that corresponded to the 3rd and 4th presentation of this sentence. If the sentence was presented consecutively, we eliminated the trials that corresponded to the 2nd and 4th presentation of this sentence.

### Speech Processing

The speech signals were first downsampled from 44.1 kHz to 8 kHz and further processed through a model that reproduces the cochlear filter bank, that is, decomposes the auditory input into 128 channels of different frequency bands spanning the range from 180 Hz to 7246 Hz (NSL Toolbox). The speech envelope from each sub-band was extracted by applying the filter bank (Chi et al., 2005), resulting in signals downsampled to 1 kHz. From each sub-band, we extracted the envelope by taking the absolute value of each time-frequency point and raising it to the power of 0.6 (Biesmans et al., 2017). The resulting 128 sub-band envelopes were averaged to obtain one single envelope, and further filtered using a minimum order bandpass FIR filter with Kaiser window between 3 and 8 Hz, with 60 dB attenuation at 2 Hz and 9 Hz respectively (Figure 2).

**Figure 2:**
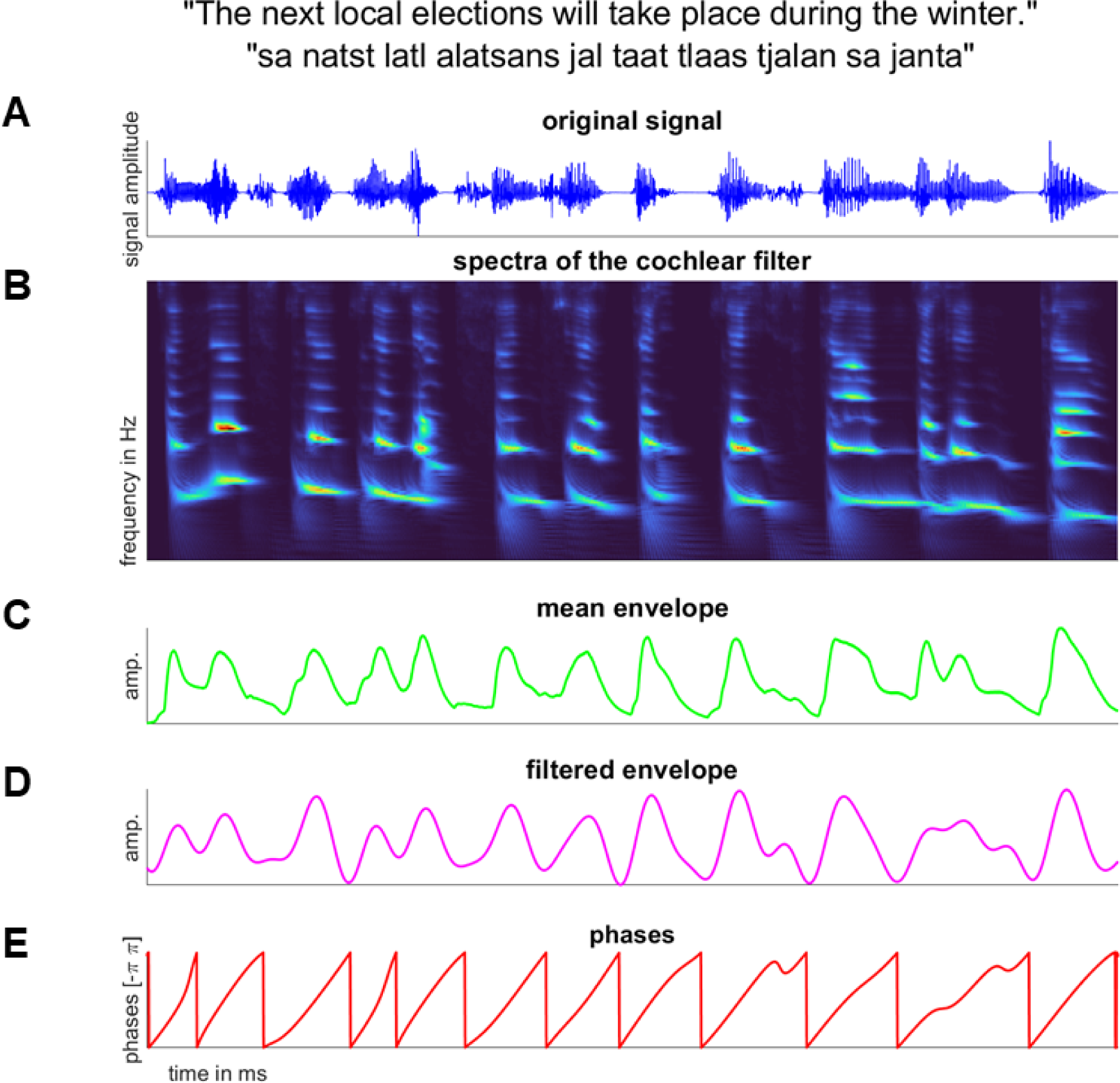
The preprocessing of the speech signals presented in the experiment. A) The sound pressure wave of one *saltanaj* resynthesized English sentence (original: “The next local elections will take place in the winter”). B) The auditory spectra of the signal after cochlear filtering. C) The mean envelope of the sub-bands extracted from the cochlear filter. D) The filtered envelope in the theta frequency (3–8 Hz). E) The phases of the Hilbert transformed signal.

### Phase Locking Value

We calculated the analytical signal for the filtered EEG via Hilbert transform at each channel and trial to extract the phases. We applied the same procedure to the filtered speech envelopes. To avoid the phase locking caused by auditory evoked responses, we removed the first 500 milliseconds of the trials from EEG and computed our measures between EEG and speech signals matched in duration (Zoefel and VanRullen, 2016). We computed PLV between the filtered EEG and speech envelopes of participant *p* at trial *t* and channel *c* as follows:

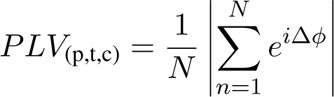

where Δ*ϕ* is the unwrapped phase differences between the Hilbert-transformed signals for each time point *n* (where *N* is the total number of time points) and *| · |* stands for the absolute value operator (Lachaux et al., 1999). PLV varies between 0 and 1: a value of 1 means full coupling, values close to 1 mean that the phase differences are extremely small, and 0 means that the phase difference between the signals varies widely, showing no coupling.

### Comparison against chance

To establish if the PLV between speech envelope and EEG reflect true phase locking, we computed PLV between speech envelopes and surrogate data, obtained by shuffling the EEG signals along the time dimension 1000 times, thus creating white-noise-like signals. We calculated the p-value by comparing the observed PLV against the PLVs between speech envelopes and shuffled EEG.

### Within group comparisons

To compare the differences in PLV across the languages, we classified PLVs from each experiment according to three conditions: stress (English), syllable (Spanish), and mora (Japanese). For each condition pair comparison (e.g., stress vs mora), we calculated the differences between the mean PLVs. As well as these differences, we obtained surrogate differences by shuffling the concatenated matrices of condition pairs across trials, and reassigning them into surrogate matrices 1000 times. We divided these surrogate matrices into two pseudo-conditions and calculated the differences in mean PLVs between these pseudo-conditions. The p-value was calculated by comparing the differences between condition pairs against the surrogate differences.

### Between group comparisons

We aimed at comparing differences in PLV to different languages between native speakers of English and Spanish. We conducted the permutation test procedure as the within group comparisons, this time, shuffling the concatenated matrices across participants instead of trials. We then divided the surrogate matrices into two pseudo-groups.

## Results

PLVs from all conditions in both experiments were significantly higher than chance values, i.e., we calculated PLV between speech envelopes and shuffled, white-noise-like EEG signals (p *<* 0.001).

### Experiment 1: Native speakers of English

PLVs for mora-timed sentences (M = 0.218, SD = 0.0056) were significantly higher than for syllable- (M = 0.213, SD = 0.0055) and for stress-timed sentences (M = 0.209, SD = 0.0062) (both p *<* 0.05). PLVs for syllable-timed sentences were also significantly higher than for stress-timed sentences (p *<* 0.05) (Figure 3).

**Figure 3:**
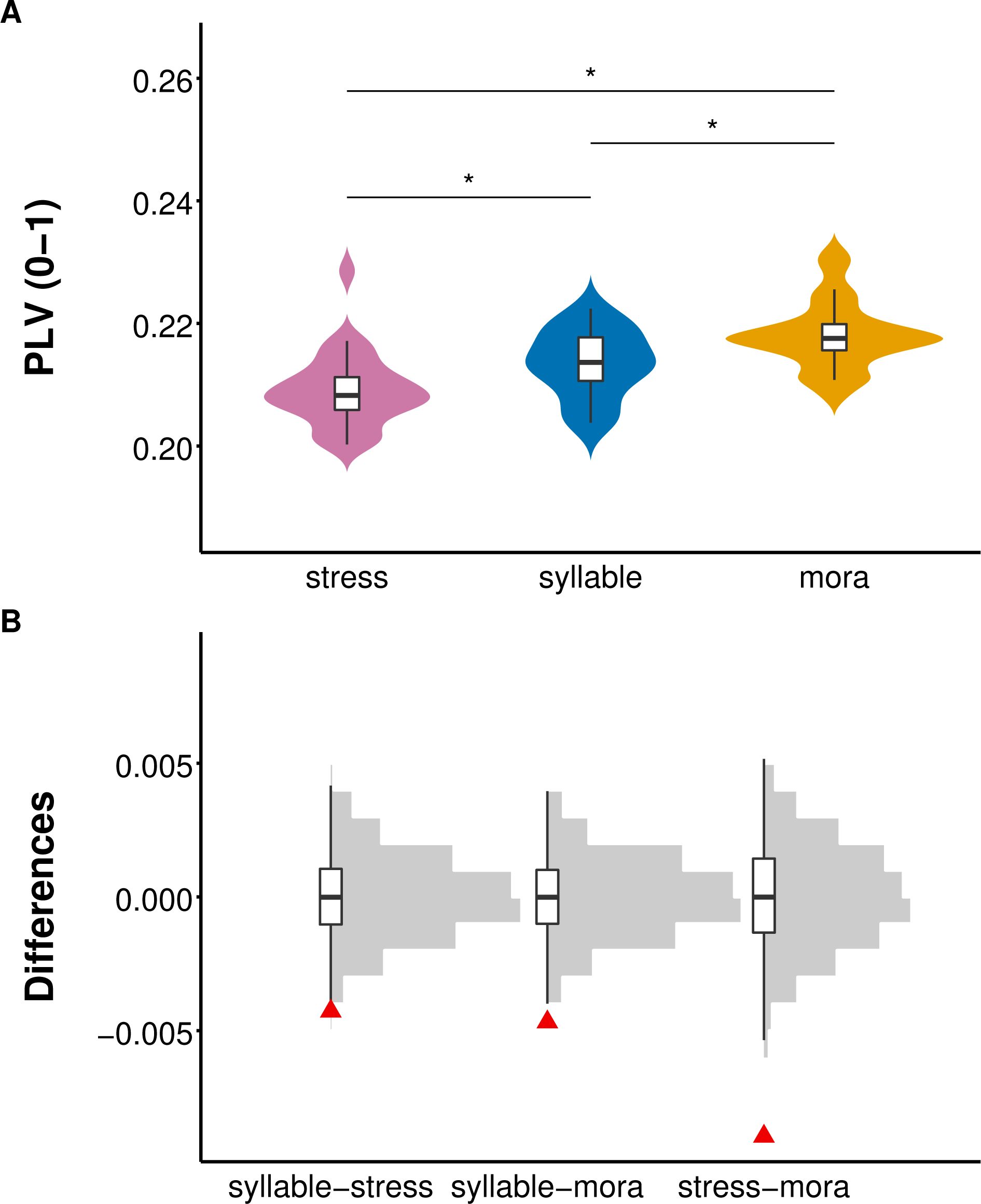
Experiment 1, native speakers of English. A) Phase locking value. Boxplots show the medians in black lines, first and third quartiles in the box edges, and minimum and maximum values of PLV at the whiskers for each condition. B) Surrogate distributions of differences (grey histogram) in each condition pair, compared to the measured differences (red triangle). Boxplots show the medians in black lines, first and third quartiles in the box edges, and minimum and maximum values of differences between pseudo-condition pairs at the whiskers for each condition.

### Experiment 2: Native speakers of Spanish

PLVs for mora-timed sentences (M = 0.217, SD = 0.0048) were significantly higher than for syllable- (M = 0.213, SD = 0.0049) and for stress-timed sentences (M = 0.211, SD = 0.0056) (both p *<* 0.05). PLVs for syllable-timed sentences did not differ significantly from those for stress-timed sentences (p = 0.061) (Figure 4).

**Figure 4:**
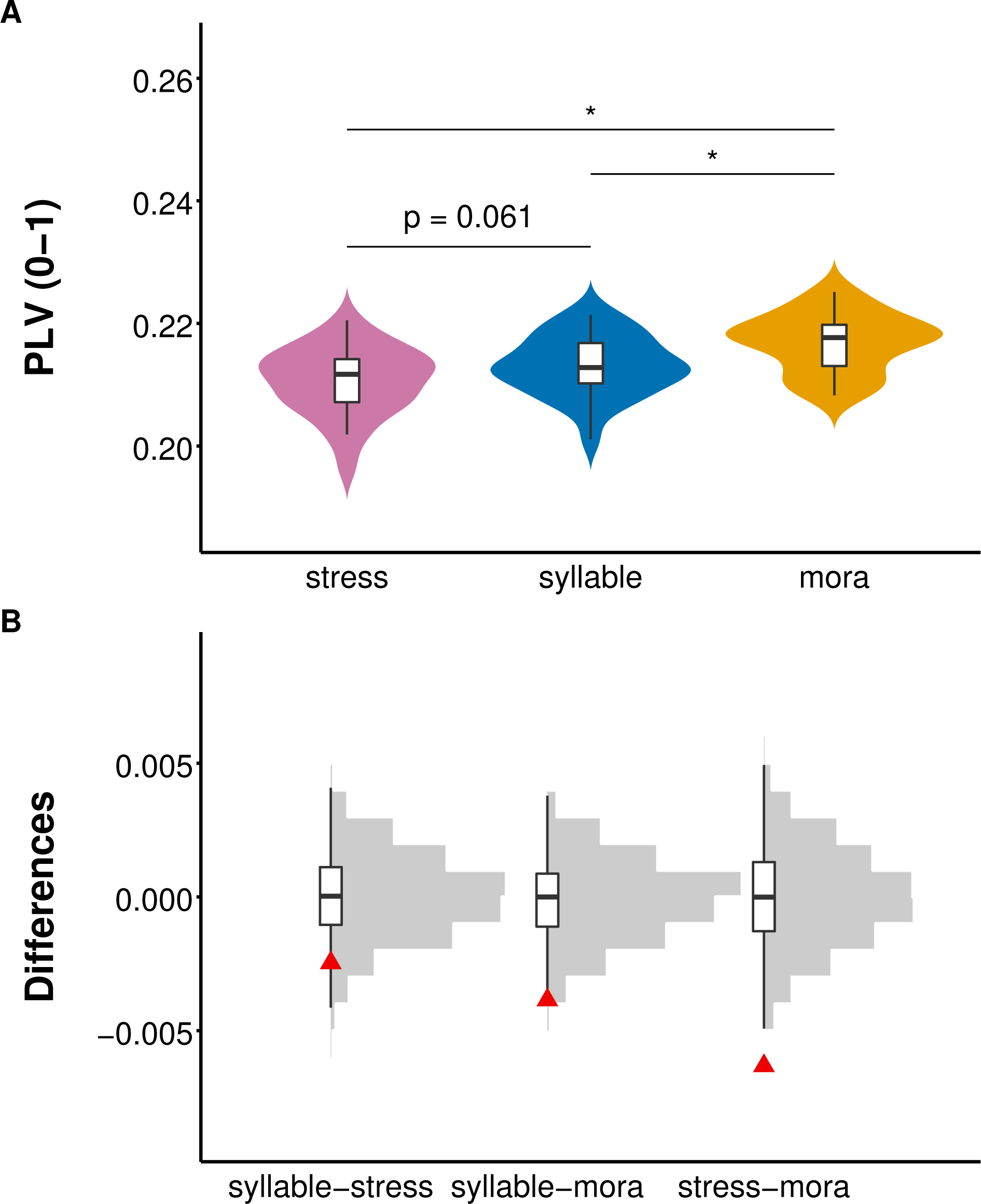
Experiment 2, native speakers of Spanish. A) Phase locking value. Boxplots show the medians in black lines, first and third quartiles in the box edges, and minimum and maximum values of PLV at the whiskers for each condition. B) Surrogate distributions of differences (grey histogram) in each condition pair, compared to the measured differences (red triangle). Boxplots show the medians in black lines, first and third quartiles in the box edges, and minimum and maximum values of differences between pseudo-condition pairs at the whiskers for each condition.

### Between-group comparisons

Permutation tests resulted in no significant difference between native speakers of English and Spanish in PLV for stress-timed sentences (p = 0.211), syllable-timed sentences (p = 0.416), or mora-timed sentences (p = 0.209) (Figure 5).

**Figure 5:**
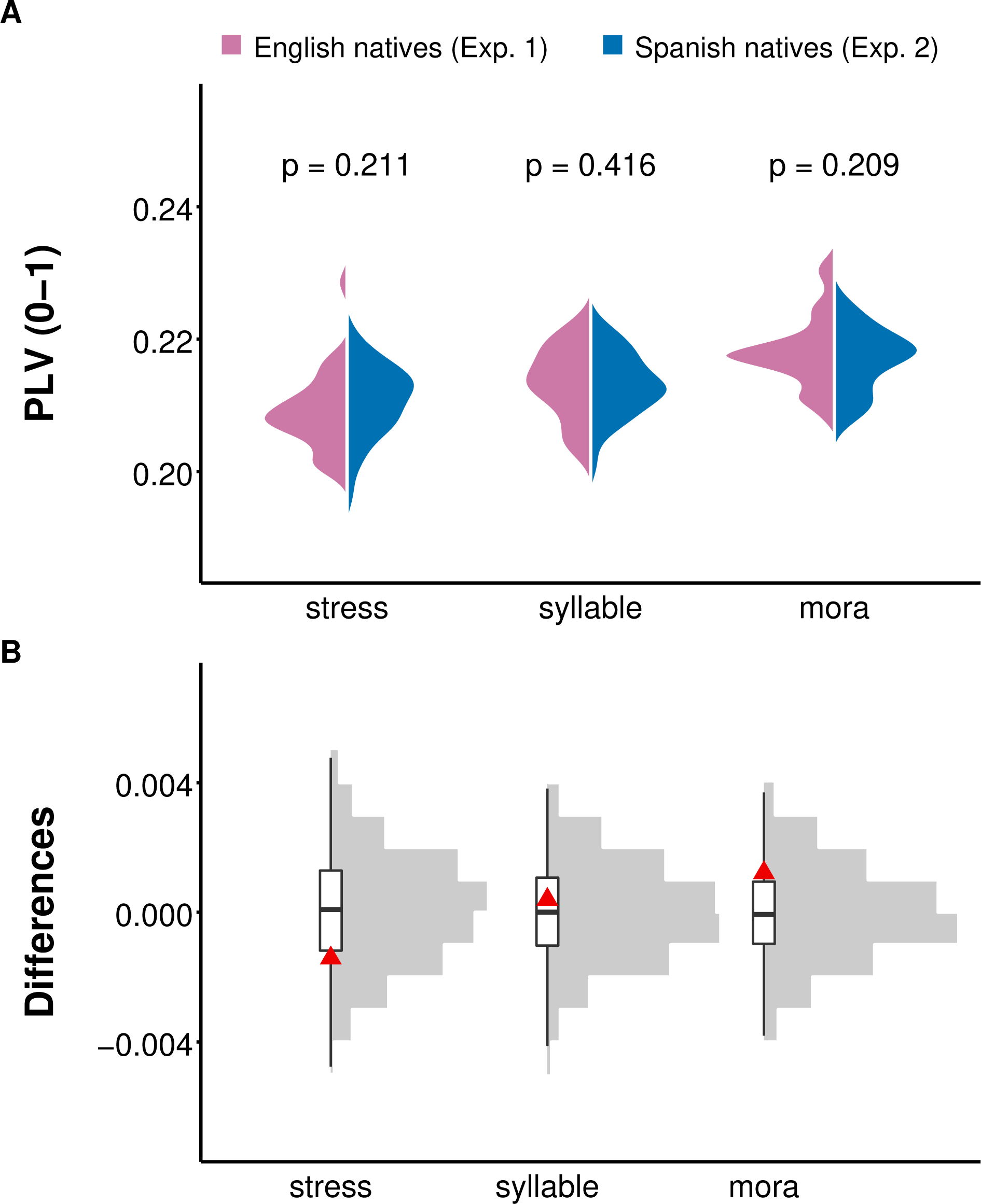
Comparison of PLV for each language across groups. A) PLVs for the sentences from the same language were compared between native speakers of English and those of Spanish. B) Surrogate distributions of mean differences (grey histogram) in each native speaker group pair, compared to the measured mean differences (red triangle). Boxplots show the medians in black lines, first and third quartiles in the box edges, and minimum and maximum values of differences between pseudo-condition pairs at the whiskers for each condition.

### Time-Frequency Analyses of the Experimental Sentences

Previous research suggests that the periodicity in time series like speech can be observed using Fourier transform by detecting the dominant beats (Ravignani and Norton, 2017). Since speech is not a completely periodic signal, and in order to overcome the violation of stationarity assumption of the Fourier transform, we used wavelet transformation (Cohen, 2014). We used wavelet cycles logarithmically increasing from 10 to 12 and obtained a frequency resolution of 0.25 Hz between 3 and 8 Hz. We averaged the time-frequency matrices across sentences and compared the results for each language using ANOVA, F(2, 60) = 7, p *<* 0.05. We observed that the power in languages show a similar pattern to that of PLV to languages, with an increasing trend from English, Spanish, and Japanese (Figure 6).

**Figure 6:**
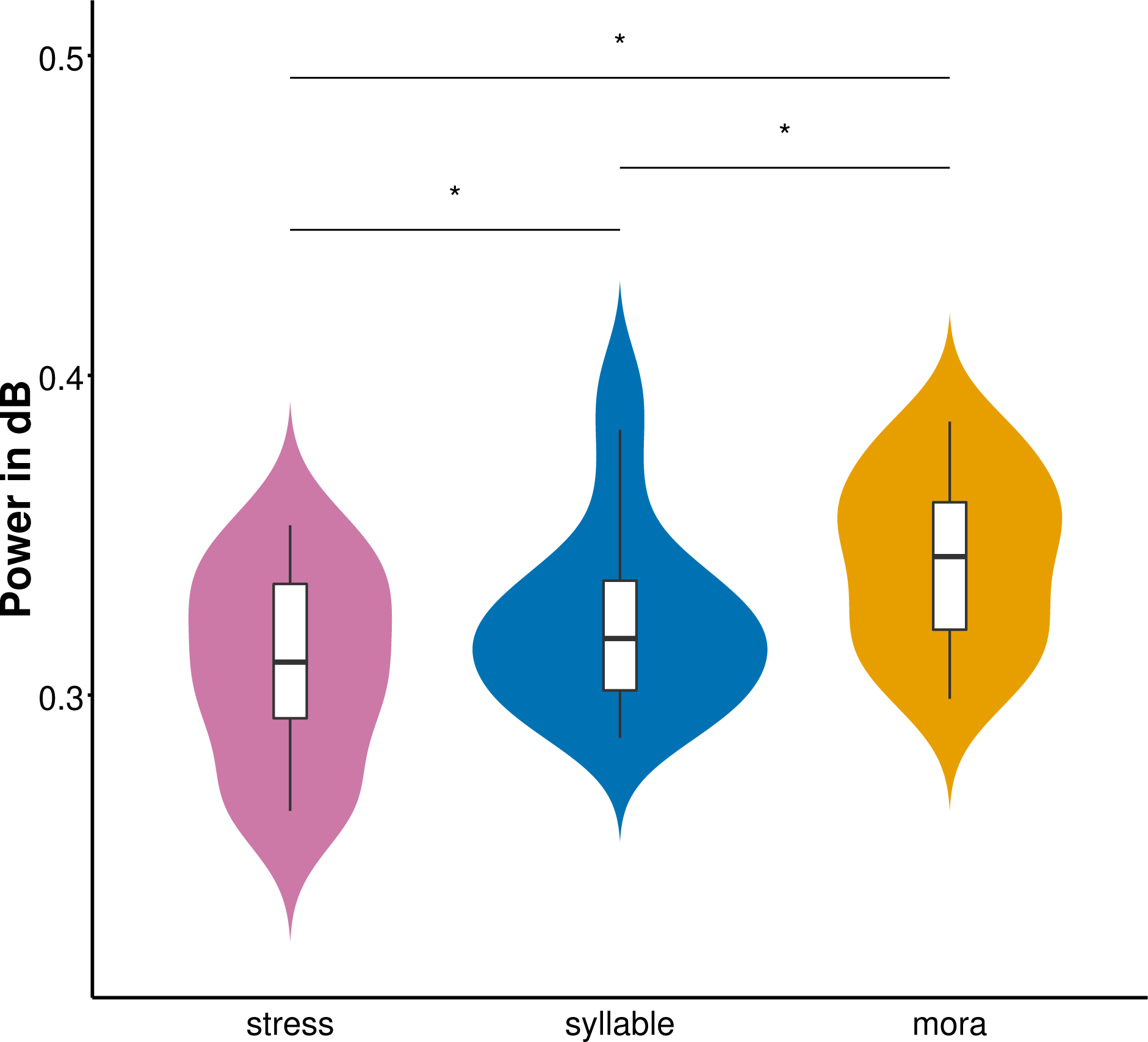
Distribution of power in theta frequency range averaged across the sentences. Boxplots show the medians in black lines, first and third quartiles in the box edges, and minimum and maximum values of power at the whiskers for each condition.

## Discussion

We conducted two experiments where native speakers of English (Experiment 1) and native speakers of Spanish (Experiment 2) listened to resynthesized sentences from English, Spanish, and Japanese. We measured phase locking value (PLV) in the theta frequency range (3–8 Hz). The common pattern in both groups indicated the lowest PLV for English, intermediate for Spanish and highest for Japanese. The comparison of PLV to each language across both groups showed no differences between native listeners of English and of Spanish. Our results indicate that neural entrainment to speech in the theta frequency range is not native-language-specific, but it is not identical for all languages.

The main goal of the present study was to test two alternative hypotheses concerning neural entrainment at the theta band: 1) participants would show stronger neural entrainment to the syllables of the resynthesized sentences from their native language, in accordance with the behavioral evidence reviewed in the introduction; 2) neural entrainment would reflect the rhythmic regularity of syllables in each language, regardless of participants’ native language and in the absence of language comprehension. According to the first hypothesis, native listeners of English should show higher PLV for English than for Spanish or Japanese and, conversely, native listeners of Spanish should show higher PLV for Spanish than for English or Japanese. According to the second hypothesis, both groups should show the same pattern: lower PLV for English, intermediate for Spanish and higher for Japanese. Our results support the second hypothesis and show no hint of better entrainment to the native language. These results partially support previous claims that neural entrainment at the theta level is not modulated by listeners’ native language, but challenge the assumption that neural entrainment to all languages share the same rhythmicity at the theta range. As mentioned in the introduction, Ding et al. (2016) and Etard and Reichenbach (2019) argued against cross-language differences in neural entrainment in the theta range. Concerning Etard and Reichenbach (2019)’s results, the lack of such differences between Dutch and English is consistent with our results, as both are stress-timed languages with a very similar rhythm. The findings of Ding et al. (2016) reporting comparable results between native speakers of English and those of Mandarin when listening to Mandarin sentences must be interpreted with caution: the lack of cross-linguistic differences might be explained by the removal of language-specific rhythmic information since the duration of all syllables was fixed to 250 milliseconds.

The assumption of a universal rhythm in speech is fundamentally based on corpora analyses (Ding et al., 2017; Varnet et al., 2017). We argue that such studies may have lacked sensitivity to detect differences in linguistic rhythm across languages. Ding et al. (2017) found a salient peak at around 5 Hz across eight different languages in their speech corpora. This peak rate was interpreted as a critical feature of speech, such as luminance in visual perception (Poeppel and Assaneo, 2020). Although the authors carefully included in their corpora a variety of speaking styles (interviews, telephone conversations, audio-books, etc.), the variability in terms of linguistic rhythm was restricted. The languages in Ding et al. (2017)’s corpora analyses were Mandarin, French, American English, German, Dutch, Danish, Norwegian, and Swedish. Except for Mandarin and French, these languages are classified as stress-timed. Furthermore, sample sizes of the different languages were not equivalent: for instance, English was analyzed over hundreds of minutes of recordings while others such as Chinese were analyzed with under one hour of recording. It is also worth noting that although all languages peaked between 4–5 Hz, the authors reported variability between languages (“… between 4.3 and 5.4 Hz for all tested speech materials” p. 183), but such variation was not further analyzed. Varnet et al. (2017) used a more varied sample in terms of linguistic rhythm: Dutch, English, Polish, French, Spanish, Turkish, Zulu, Basque, Marathi and Japanese. While this was not the primary goal of their research (and was therefore not further elaborated on), their analyses pointed to a difference in the peak rate between the languages from the syllable- and stress-timed rhythms. The analysis of the power modulation spectra in the theta range of our sentences yielded differences between the three languages (Figure 6). Our results are consistent with the notion of increased regularity in linguistic rhythm in English, Spanish and Japanese, reflecting the decrease in syllabic complexity present in each language (see Langus et al. 2017 for an equivalent proposal in the linguistic domain). We acknowledge the small sample of languages and sentences used in our own work, limiting the generalizability of our results. Future studies should analyze whether the differences in terms of linguistic rhythm and power at the theta range we report here are replicated with larger corpora testing a wide range of languages sufficiently varying in their linguistic rhythm.

The results of our study do not challenge the idea that neural entrainment in theta band depends on a universal syllabic rate in the speech stream (Ding et al., 2016, 2017; Assaneo and Poeppel, 2018; Poeppel and Assaneo, 2020). Rather, we extend this idea by providing evidence that neural entrainment in the theta range is modulated by the variation in syllabic complexity across languages. In our experiments, theta oscillations entrained best to resynthesized sentences with reduced variation in syllabic complexity (mora-timed). In this regard, it is worth noticing that the syllable types in Japanese (V, CV and CVC) are also the most common in the world languages (Greenberg, 1999).

One additional novelty of our research is to measure neural entrainment to *saltanaj* resynthesized sentences to eliminate the confounding effects of familiarity with stimuli (whether native or non-native), and critically, the comprehension of participants’ native language. The validity of this resynthesis method in speech processing was previously established in psycholinguistic experiments. It was found that native French adults can discriminate between resynthesized English and Japanese (Ramus and Mehler, 1999), and French-learning newborns can discriminate between resynthesized Japanese and Dutch (Ramus, 2002). The use of resynthesis to replace different phonemes is not an uncommon practice in psycholinguistic experiments (Christophe et al., 2003; Herment-Dujardin and Hirst, 2002). By using resynthesized sentences, we managed to reduce the phonological differences in our stimuli and concentrate our results to the differences in syllabic complexity across languages.

The results of our work open two questions. First, what are the processing consequences of better bottom-up entrainment for certain languages? Second, what is the origin of the cross-language differences observed in the behavioral studies reported in the introduction? Neural entrainment to speech is the product of top-down and bottom-up factors (Rimmele et al., 2018; Assaneo et al., 2019). It is associated with increased comprehension of sentences (Peelle et al., 2013), with certain studies showing that intelligibility is compromised when brain rhythms cannot align with acoustic ones. Bosker and Ghitza (2018) presented participants with lists of stimuli (digits) compressed at 3, 6, and 15 Hz, and showed that intelligibility performances dropped at 15 Hz (outside the theta range). Assuming that the speech processing system is optimally fit for language comprehension, we propose that less efficient bottom-up processing may be compensated by stronger top-down processing, either by increased reliance on linguistic knowledge, or by recruiting other types of domain-general mechanisms, such as attention. Such stronger reliance should be detected in situations when comprehension is challenging, as native speech processing in adults might be efficient enough to compensate for the slight processing advantages provided by linguistic rhythm in the bottom-up component (Blanco-Elorrieta et al., 2020). Concerning the second question, in this study we originally attempted to explain the psycholinguistic evidence suggesting that rhythmic properties of listeners’ native language impacted the way the speech signal was initially segmented (Mehler et al., 1981; Cutler et al., 1986). Behavioral experiments following Mehler et al. (1981) and Cutler et al. (1986)’s original results showing contrasting speech segmentation between French and English listeners have been replicated in a variety of languages and laboratories (Catalan and Spanish: Sebastián-Gallés et al. 1992; English and Spanish: Bradley et al. 1993; Japanese: Otake et al. 1993; Dutch: Zwitserlood et al. 1993; French: Dumay et al. 2002, among others). In these studies, cross-language behavioral differences were proposed to reflect low-level parsing of the signal, showing plasticity to native phonology (Mehler et al., 1988; Ramus, 2002). If entrainment at the theta band reflects low-level linguistic chunking of the speech signal (Boucher et al., 2019), we should have observed differences between speakers of different languages, namely better entrainment to stimuli following the properties of the native language. However, our results failed to support this hypothesis. Unfortunately, our results shed no light on the origin of the cross-language differences reported in such studies reported, except for the fact that they do not reflect initial speech segmentation processing.

To conclude, we provide evidence of language-specific modulation in neural entrainment at the theta range, reflecting properties of the linguistic rhythm regardless of listeners’ native language. These findings closely match the differences we observed in power of the speech signal at the theta range.

